# Hypothesis Testing Governs an Efficiency-Flexibility Trade-off in Strategic Motor Learning

**DOI:** 10.64898/2025.11.29.691289

**Authors:** Wei Ding, Anjuli Niyogi, Jonathan S. Tsay

## Abstract

It remains unknown how people discover an effective movement strategy when the environment changes (e.g., when adapting to a new computer trackpad). We propose that strategic adaptation operates through hypothesis testing: learners generate candidate hypotheses, discard those inconsistent with feedback, and iteratively refine their actions through practice. A core prediction of this account is an efficiency–flexibility trade-off. In constrained environments, where few hypotheses are viable, learning slows as people eliminate competing hypotheses but supports broader generalization. In unconstrained environments, where many hypotheses are viable, learning accelerates as learners adopt one of many expedient hypotheses but yields poorer generalization. In two reaching experiments (N = 560), we varied the arrangement of target positions to manipulate how tightly the hypothesis space was constrained. As predicted, the constrained group learned more slowly but generalized more—an efficiency– flexibility trade-off that highlights hypothesis testing as a novel process governing human motor learning.

## Introduction

Successful goal-directed action depends on multiple learning processes (Shadmehr et al., 2010). Among these, strategic motor adaptation is especially critical for efficiently adjusting our movements in response to changes in the body and in the environment (McDougle et al., 2016; Taylor & Ivry, 2011). For example, when switching to a new ping-pong paddle, players often *deliberately* adjust their swing angle to compensate for changes in bounce, rapidly forming an *explicit strategy* to maintain precision and control.

Traditionally, strategic (or explicit) adaptation has been framed as a gradual process of error minimization: when the environment changes, learners incrementally reduce their motor errors to restore success (Taylor & Ivry, 2011). This view aligns with the long-standing assumption that motor learning follows a smooth power law—treated for decades as a foundational principle of the field (Snoddy, 1926; Crossman, 1959; Logan, 1988). Support for this perspective comes in part from the classic visuomotor rotation task, which perturbs the mapping between visual feedback and movement direction. Group-averaged performance in counteracting the perturbation follows a smooth, power-law trajectory (Coltman et al., 2021; Haith et al., 2015; Smith et al., 2006; Taylor & Ivry, 2011), a signature taken as evidence for gradual error reduction (Krakauer et al., 2019). Any behavioral variability around the smooth power-law curve is attributed to small amounts of noise (5°-10°) from the sensorimotor system (Albert & Shadmehr, 2018).

But individual learning curves reveal a very different story. Rather than gradually converging on the solution, learners often engage in rich, exploratory behavior that is anything but gradual, with variability that is anything but minimal (Townsend et al., 2023; see review: Tsay et al., 2024b). Furthermore, movement errors cluster into multiple modes, often far from the solution (Chen et al., 2025; McDougle et al., 2019)—a pattern incompatible with gradual error reduction, which predicts motor errors confined between baseline and solution states. Consequently, by averaging across participants, group-level analyses mask the variability that defines strategic motor adaptation (Abe & Sternad, 2013; Sternad, 2018; Sternad et al., 2011).

Inspired by individual learning data, we have conjectured a novel framework for strategic adaptation: hypothesis testing (Tsay et al., 2024b). When confronted with a new visuomotor mapping—such as the altered bounce from an unfamiliar ping-pong paddle—learners generate candidate hypotheses for the environmental change (e.g., that the paddle induces an added rotation or a mirror reversal), test them against sensory feedback, discard those that fail, and iteratively refine both their actions and their hypotheses through practice (Chen et al., 2025; Niyogi et al., 2024). Indeed, hypothesis testing offers a fundamentally different view of strategic adaptation, explaining the exploratory, individual-level trajectories where learning processes unfold.

Here, we directly test a prediction unique to Hypothesis Testing: the efficiency–flexibility trade-off (Collins & Frank, 2013; Collins, 2017; Del Giudice & Crespi, 2018; Lieder & Griffiths, 2020). In constrained environments, where few hypotheses are viable, learning slows as learners iteratively eliminate competing hypotheses (inefficient) but ultimately promotes generalization (flexible), that is, the application of the learned rule in a new context. In unconstrained environments, where many hypotheses are viable, learning accelerates as learners adopt one of many expedient strategies (efficient) but ultimately compromises generalization (inflexible). Because gradual error reduction accounts do not offer a clear prediction about how environmental constraint should shape learning or generalization, demonstrating this trade-off would directly implicate hypothesis testing as a core mechanism of strategic adaptation.

In search of an efficiency–flexibility trade-off, we conducted two large-scale experiments (N = 560) using a visuomotor rotation task specifically designed to isolate strategic motor adaptation (Brudner et al., 2016). Participants were instructed to explore the workspace to align a 60° rotated cursor to the target without being told the nature of the perturbation (Figure 1A-B). Critically, we manipulated the level of environmental constraint via the spatial geometry of the training targets (Figure 1C–D). In the constrained geometry, widely spaced outer targets produced action–outcome patterns that prompted a rotational hypothesis (Figure 1E, left). In the unconstrained geometry, closely spaced inner targets allowed multiple hypotheses to generate success during training (Figure 1E, right).

**Figure 1.**
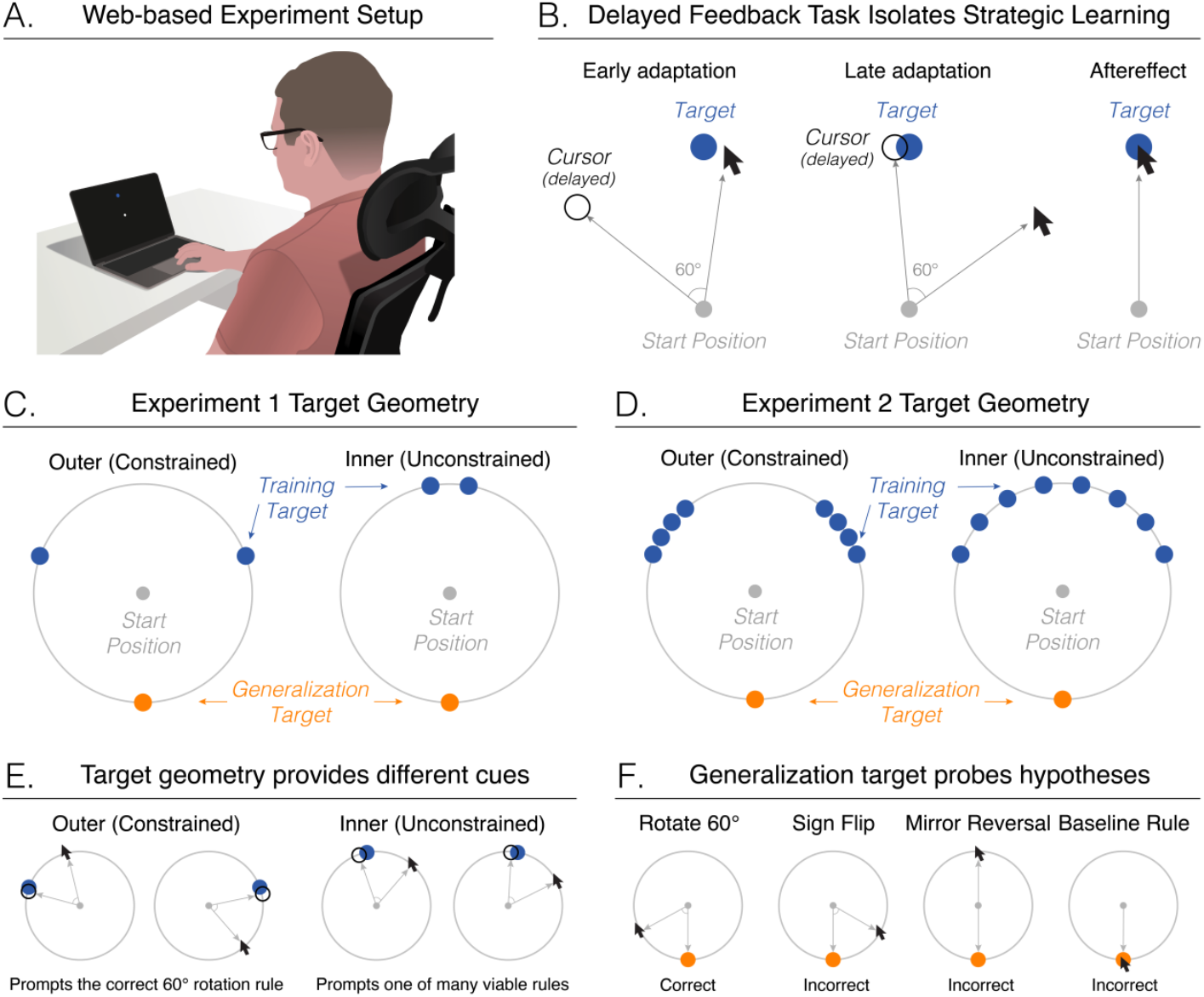
Experimental Overview. **(A)** Web-Based Experimental Setup. Participants completed a visuomotor adaptation task using their computer trackpad. **(B)** The Delayed Feedback Task isolates strategic motor adaptation. Endpoint cursor feedback, rotated by 60° (hollow black circle), was displayed 1000 ms after the hand (in screen coordinates) reached the target distance—a manipulation known to minimize implicit adaptation. Participants were instructed to explore different movements to align the rotated cursor with the target. Importantly, they were not told about the nature of the perturbation. **(C)** Experiment 1: Inner vs. outer target geometries. Participants reached to two training targets (blue circles) spaced either far apart (outer group, 140°) or close together (inner group, 20°). A single generalization target (orange circle) was positioned opposite the midpoint between the training targets. **(D)** Experiment 2: Inner vs. outer target geometries. Eight training targets were arranged either along the outer edge of the circle (outer group, 5° spacing) or more uniformly across the workspace (inner group, 20° spacing). Unlike Experiment 1, the distance between training and generalization targets was equated across groups. The gray circle was not shown to participants and is included here only for illustrative purposes. **(E)** Example movements (cursor pointer) and cursor outcome (hollow black circles) at the two training targets in Experiment 1 for the Outer- and Inner-target groups. **(F)** Possible movements at the no-feedback generalization target, each reflecting a learned action–outcome hypothesis: the correct imposed rotation (60° clockwise) or one of several incorrect rules, including the sign-flipped rotation (60° counterclockwise), a mirror reversal (180°), or a default reach-to-target response (0°).

To track how learners’ action–outcome hypotheses evolved, we interleaved a no-feedback generalization target throughout the experiment. At this target, participants could reveal whether they inferred the correct rotational rule (60° clockwise) or one of several incorrect rules, such as a sign-flipped rotation (60° counterclockwise), a mirror reversal (180°), or a default baseline response (0°) (Figure 1F). Together, these experiments directly test the predicted trade-off—slower but more generalizable learning in the outer-target groups—and evaluate hypothesis testing as a core process governing human sensorimotor learning.

## Methods

### Participants and Apparatus

We recruited 560 participants through Prolific, an online participant recruitment website (312 females, aged 25.1 ± 3.4 years, all participants self-reported being right-handed). This sample size is substantially larger than that of typical in-person motor learning studies (Tsay et al., 2021). Before beginning the study, all participants provided informed consent in accordance with policies approved by Carnegie Mellon University’s Institutional Review Board. Participation was in exchange for monetary compensation.

The experiment was created using the OnPoint platform, a package for running customized online motor learning experiments with JavaScript (Tsay et al., 2021). Participants completed the web-based experiment via an internet browser with their laptop’s trackpad (Figure 1A). Our past online studies have shown that neither the type of browser nor the type of pointing device impacts motor performance (Tsay et al., 2024a). The size and position of the visual stimuli were scaled to the individual’s monitor size. Specifically, the workspace was centered on the screen midpoint, and target eccentricity was fixed at 25% of the screen height. All other stimuli were scaled according to the same proportion of screen height. For ease of interpretation, all stimulus parameters detailed below were based on an average 13-inch computer monitor.

### General Procedure

We employed the Delayed Feedback Task to isolate strategic adaptation (Brudner et al., 2016; Tsay et al., 2023; Figure 1B). On each trial, participants positioned a white cursor (0.3 cm diameter) inside a white starting ring (0.5 cm diameter) at the center of the screen. Once the cursor entered the ring, it filled in. After holding the cursor there for 500 ms, the cursor disappeared, and a target (0.5 cm diameter) appeared on an invisible ring with a 4.5 cm radius. The target cued participants to make a rapid, straight movement to slice through it. When the hidden cursor crossed the invisible 4.5 cm ring, an auditory beep signaled movement termination. Visual cursor feedback reappeared 1000 ms after movement termination—a delay shown to minimize implicit processes underlying motor adaptation (Kitazawa et al., 1995)—and remained visible for 500 ms. Participants then returned to the start position for the next trial. To support reproducibility, full task instructions are provided in the Supplementary Materials.

### Experiment 1

Participants (N = 280; 155 females; mean age = 25.9 ± 3.8 years) were randomly assigned to one of two groups. In the inner-target (unconstrained) group (N = 140), two blue training targets were positioned close together (20° apart) along with a single generalization target positioned opposite the midpoint between the training targets (Figure 1C). In the outer-target (constrained) group (N = 140), the training targets were positioned farther apart (140° apart) with the same generalization target. To ensure the effects were not specific to particular target locations, we counterbalanced target placement across participants (Table S1).

Trials were organized into triplets (3 trials/cycle): Two reaches to blue training targets with visual feedback followed by one reach to an orange generalization target without feedback. The task comprised of 132 trials (44 reaches/target), divided into three blocks: baseline (24 trials; 8 reaches/target), perturbation (96 trials; 32 reaches/target), and aftereffect blocks (12 trials; 4 reaches/target). During the baseline block, cursor feedback was veridical—aligned with the participant’s actual movement direction—and was the only block in which visual feedback was provided for the generalization target. In the perturbation block, a 60° visuomotor rotation was applied to cursor feedback relative to the participant’s movement direction, with the direction of rotation (clockwise vs counterclockwise) counterbalanced across participants. Rotated feedback was only provided at the training target; no feedback was given at the generalization target. In the aftereffect block, no visual feedback was provided at either training or generalization targets.

### Experiment 2

All procedures in Experiment 2 were identical to those in Experiment 1, except that Experiment 1 included *two* training targets, whereas Experiment 2 included *eight*. Participants (N = 280; 156 females; mean age = 24.9 ± 3.4 years) were randomly assigned to one of two groups. In the inner-target group (N = 140), the eight training targets were spaced 20° apart (Figure 1D). In the outer-target group (N = 140), the eight training targets were spaced 5° apart. In both groups, the nearest target lies 110° from the generalization target, thereby equating this factor across inner- and outer-target conditions. To ensure the effects were not specific to particular target locations, we counterbalanced target placement across participants (Table S1). Trials were organized into triplets (3 trials/cycle). The task comprised 132 trials (11 trials/training target; 44 trials/generalization target), consisting of baseline (24 trials; 2 trials/training target, 8 trials/generalization target), perturbation (96 trials; 8 trials/training target, 32 trials/generalization target), and aftereffect blocks (12 trials; 1 trial/training target, 4 trials/generalization target).

### Data analysis

All data processing and statistical analyses were conducted in Python 3.9.7. Analyses focused on hand position in screen coordinates (i.e., the position of the computer pointer) at the point where movement amplitude reached the target radius (4.5 cm). These data were used to calculate our main dependent variable, movement angle, defined as the angular distance between the movement and target positions. Movement angles across perturbation directions (clockwise vs. counterclockwise) were standardized so that positive values always indicated changes that counteracted the perturbation (i.e., movement angles for the counterclockwise rotation groups were multiplied by –1).

We summarized movement angles across three phases: early adaptation, late adaptation, and aftereffect. Early adaptation was defined as the average change in movement angle over the first 4 cycles of the perturbation block (cycles 9–12). Late adaptation was defined as the average movement angle over the last 4 cycles of the perturbation block (cycles 37-40). Aftereffect was defined as the average movement angle over all 4 cycles of the no-feedback washout block (cycles 41-44). Because movement angle was measured in polar (angular) coordinates, mean angles were computed using circular statistics (Batschelet, 1981; Fisher, 1995), and variability was quantified using circular standard error of the mean (SEM). All key results were also replicated under linear, non-circular statistics.

To evaluate group differences between inner (unconstrained) and outer-target (constrained) groups, we conducted a 2 (Target Type: training vs. generalization) x 2 (Target Geometry: inner vs. outer) mixed-design ANOVAs on early adaptation and late adaptation phases. Target Type was a within-subjects factor and Target Geometry was a between-subjects factor. We report the F statistic, corresponding p-value, and effect sizes as partial eta squared (η^2^). Post hoc tests were conducted using Watson two-sample tests for unpaired-sample circular comparisons and paired-sample within-sample circular comparisons. An efficiency–flexibility trade-off would manifest as a significant Target Type x Target Geometry interaction, with post-hoc tests confirming that the inner-target (unconstrained) group learns faster at trained targets but generalizes more poorly to the untrained target compared to the outer-target (constrained) group.

### Identifying Clustered Differences in Learning Functions

To identify differences between inner- and outer-target geometries without imposing arbitrary phase boundaries (early vs late), we applied a cluster-based permutation test to movement angles (Breska & Ivry, 2020; Tsay et al., 2020; Wang et al., 2024). Training and generalization trials were analyzed separately. Statistical comparisons were performed using the Watson two-sample test (U^2^-values), which yields a p-value assessing whether two circular distributions differ significantly. Movement data were analyzed in two-cycle units. Units with a p <.05 were marked as candidates for clustering, and clusters were defined as two or more adjacent significant units. The cluster-level test statistic was defined as the sum of U^2^-values across constituent units. Cluster significance was assessed against a null distribution generated from 5,000 permutations in which group labels were shuffled. For each permutation, the clustering procedure was repeated and the largest cluster sum recorded. Clusters were deemed reliable if their summed statistic exceeded the 95th percentile of the null distribution (α =.05), thereby controlling the false positive rate. Effect sizes for each significant cluster were reported as (1) the absolute circular mean difference (Δμ, in degrees) between groups, (2) circular Cohen’s d and (3) the permutation-based cluster p-value (Mardia & Jupp, 2000; Pewsey et al., 2013).

### Characterizing Multimodal Motor Error Distributions

To characterize the multimodal structure of participants’ exploratory behavior, we fit a mixture of von Mises distributions to the movement angle data using an Expectation–Maximization (EM) algorithm (Dempster et al., 1977; Banerjee et al., 2005). EM iterations were capped at 1,000 steps with a convergence tolerance of 1×10^-8^. The optimal number of mixture components (k=2–10) was determined via the Bayesian Information Criterion (BIC; Schwarz, 1978). For each k, the model was fit with 5,000 random initializations. For each component, we estimated three parameters: weight (proportion of movement angles assigned to that component), mean (degrees), and concentration parameter κ (the circular analogue of precision). We imposed four constraints to prevent degenerate model fits: (1) a minimum weight of 0.1% per component, (2) a minimum κ of 4.0, (3) a uniform distribution component to capture random responses with its weight constrained to not exceed 20%, and (4) a minimum angular separation of 25° between peaks. Additionally, to assess whether motor errors were distributed differently between the inner-target and outer-target groups, we computed Jensen–Shannon Divergence (JSD; Lin, 2002). A significant JSD (p < 0.05) would indicate that the groups produced distinct underlying distributions.

### Characterizing the Effect of Target Geometry as a Continuum

To assess whether learning varied systematically across the four target geometries in a continuous manner, we conducted mixed-effects ANOVA in which Target Geometry was encoded along a hypothesized continuum (2 outer → 8 outer → 8 inner → 2 inner). We ran these analyses separately for training and generalization trials. Evidence for a continuum would appear as a significant beta coefficient across geometries.

### Identifying Subgroups of Learners

We sought to identify distinct subgroups of learners using a two-step procedure. First, because studies of strategic adaptation frequently include participants who show little or no learning (Cisneros et al., 2024; Tsay et al., 2023; Jang et al., 2023), we removed ‘non-learners’ prior to clustering. Specifically, we applied a one-sample V-test at the subject level to late-adaptation angles (μ_0_ = 60°, p <.05) and a Watson two-sample test between the late-adaptation and aftereffect angle distributions (p <.05), yielding a sample of 247 out of 280 participants in Experiment 1 (88% learners) and 229 out of 280 participants in Experiment 2 (82% learners). Second, to identify distinct types of learners within this filtered learner sample, we applied Soft Dynamic Time Warping (Soft-DTW; γ = 0.1; Cuturi & Blondel, 2017) to each participant’s learning function. Ward’s hierarchical clustering was then performed on the resulting distance matrix. The optimal number of clusters was selected using both Silhouette Scores (Rousseeuw, 1987) and Within-Cluster Sum of Squares (WCSS; McQueen, 1967). Finally, a chi-square test evaluated whether cluster frequencies differed across target-geometry conditions.

### Data and code availability statement

All data and code are available on the Open Science Framework (OSF) via the OpenMotor repository (https://osf.io/aknqj/).

## Results

### Experiment 1: Uncovering an Efficiency-Flexibility Trade-off in Strategic Motor Adaptation

We tested for an efficiency–flexibility trade-off in strategic motor adaptation—a behavioral signature uniquely predicted by the Hypothesis Testing framework (N = 280). Specifically, we employed a visuomotor rotation task designed to isolate strategic processes (Figure 1A-B; Brudner et al., 2016): After a baseline block with veridical feedback, participants reached toward two interleaved training targets while receiving cursor feedback rotated 60° relative to their movement direction. Endpoint cursor feedback was delayed by 1000 ms—a manipulation known to minimize implicit adaptation—ensuring that learning performance primarily reflected deliberate, strategic (explicit) processes. Participants were not informed about the nature of the perturbation; instead, they were instructed to freely explore the workspace to learn the visuomotor mapping and align the perturbed cursor with the target.

To vary the level of environmental constraint, we manipulated the spatial separation between the two training targets (Figure 1C): In the outer-target group (constrained environment), the wide target separation provided strong geometric cues that the perturbation was a rotation, narrowing the set of viable hypotheses (Figure 1E, left). In contrast, the inner-target group (unconstrained environment) offered weaker, more ambiguous spatial cues, yielding a broader hypothesis space in which multiple hypotheses remained viable (Figure 1E, right). While the gradual error reduction framework offered no clear prediction about how environmental constraints shape learning and generalization, the Hypothesis Testing framework again predicted an efficiency–flexibility trade-off: the outer-target group should learn slower yet generalize more to novel targets.

Following a baseline phase to familiarize participants with the visuomotor rotation task, participants were exposed to the rotated visual feedback. Both group average learning trajectories showed a canonical, gradual learning curve at the two training targets: movements starting at the baseline target and progressing toward the 60° solution. Traditionally, this pattern has been taken to support the gradual error reduction framework (Figure 2A).

**Figure 2.**
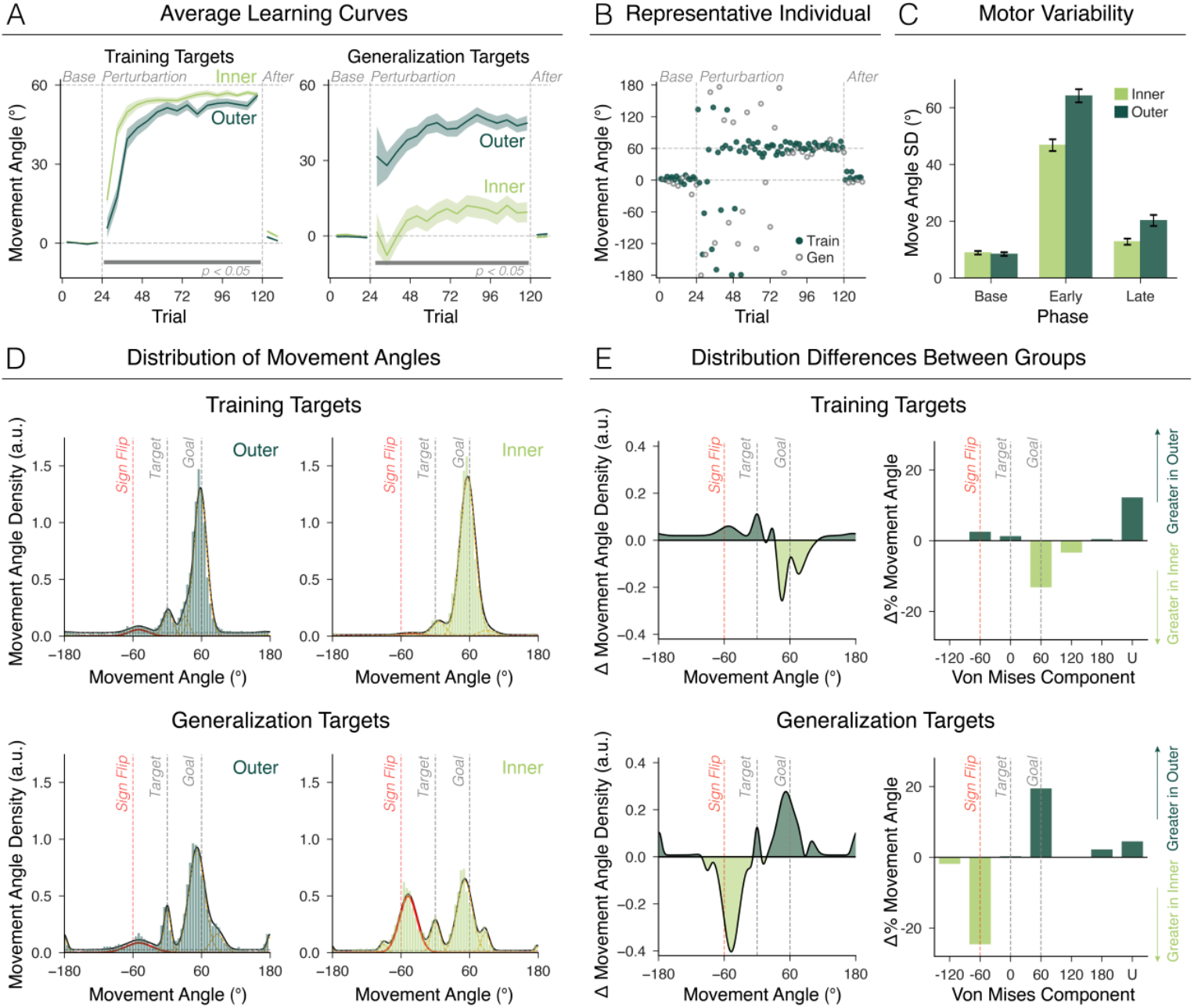
Uncovering an Efficiency–Flexibility Trade-off in Strategic Motor Adaptation. **(A)** Learning curves for training **(left)** and generalization targets **(right)** in Experiment 1. Mean movement angle (°) is plotted for the outer-target (dark green) and inner-target (light green) groups; shaded regions denote SEM. Vertical dashed lines indicate perturbation onset and offset. Horizontal gray bars mark significant group differences identified via cluster-based permutation tests (p <.05). **(B)** Representative learner from the outer-target group. Filled green dots denote movements to training targets; open grey dots denote movements to generalization targets. **(C)** Within-participant standard deviation of movement angles (motor variability) at the training target during baseline, early adaptation, and late adaptation. Error bars reflect SEM. **(D)** Movement angle distributions for training (top) and generalization (bottom) targets (density). Solid black lines show von Mises mixture fits; dashed red lines mark individual mixture components; histograms are rescaled to match the density range. Vertical dotted lines denote the training target (0°) and the required rotation solution (60°). Sign flips (–60°)—correct-magnitude reaches in the opposite/incorrect direction—are highlighted in red for illustration. **(E)** Distributional differences (density) between inner- and outer-target groups for the training and generalization targets **(left)**. Negative values indicate greater density in the inner-target group; positive values indicate greater density in the outer-target group. Group differences are corroborated by von Mises component weights (60° bins) and the uniform (U) component **(right)**.

However, a striking dissociation emerged between groups: The inner-target (unconstrained) group learned more rapidly at the training targets (Figure 2A, top) but exhibited a marked failure to generalize at the untrained target (Figure 2A, bottom). In contrast, the outer-target (constrained) group learned more slowly at training targets but generalized more flexibly to the untrained target. Together, these results uncovered an efficiency–flexibility trade-off—a pattern uniquely predicted by the hypothesis testing framework.

This efficiency–flexibility trade-off was verified statistically. Across the whole perturbation block, the inner-target group consistently outperformed the outer-target group at the training locations (cluster-based permutation test: cycles 9 – 40, Δμ = 6.4°, *d*_*circ*_ = 0.1, *p*_*perm*_ <.001) (Figure 2A, left): During early adaptation, the inner-target group adapted nearly twice that of the outer-target group (Inner: 43.1°± 2.4°; Outer: 28.5°±3.2°; p<.001), demonstrating more efficient learning under the less constrained environment. By late adaptation, both groups had converged on the 60° solution, nullifying the visuomotor rotation (Inner: 56.9° ± 1.1; Outer: 54.4° ± 1.9; *p* =.035). A one-sample V-test confirmed that movement angles during the aftereffect phase were clustered around 0° in both groups (*p* <.001), confirming that learning in both groups reflect strategic, rather than implicit, learning processes.

The pattern of group performances reversed in the generalization probes (Figure 2A, right) (interaction between Target Geometry x Target Type: *F* = 41.9, *p* <.001, *η*^2^ = 0.13): participants trained with inner-targets showed poor generalization compared to those trained in the outer-target group (cluster-based permutation test: cycles 9 – 40, Δμ = −35.1°, *d*_*circ*_ = −0.6, *p*_*perm*_<.001). Although neither group achieved full generalization (60°), the outer-target group exhibited a persistent advantage (late adaptation: Inner: 10.0° ± 4.2; Outer: 43.4° ± 3.6; p<.001), demonstrating that a more constrained environment yielded greater generalization despite less efficient learning.

In stark contrast to the smooth group-averaged curves, individual learning trajectories revealed pronounced exploratory behavior (Figure 2B; additional participants shown in Figure 4E). Motor variability during early adaptation was 46.8° ± 2.1° in the inner-target group and 64.2° ± 2.3° in the outer-target group (Figure 2C). These values far exceeded baseline variability, which presumably reflected sensorimotor noise (main effect of Phase: *F* = 856.6, *p* <.001, *η*^2^ = 0.76). Variability then declined over training (late adaptation SD; Inner: 12.8° ± 1.1, Outer: 20.3° ± 2.0). This temporal profile—large early variability followed by convergence on the solution—was incompatible with the gradual error reduction account, which assumed minimal (5° - 10°) and stationary sensorimotor noise. Instead, it aligned with learners testing multiple hypotheses early in learning and gradually settling on an effective strategy as sensory evidence accumulated.

To identify which action–outcome hypotheses participants explored, we analyzed movement-angle distributions in the early exploratory phase. Movements to the training targets were strikingly multimodal (Figure 2D, top; Best BIC, inner = −8563.3 with 6 modes; outer = −5369.6 with 5 modes). Beyond reaches to the optimal solution (60°), three prominent peaks emerged: sign flips (correct magnitude, wrong direction; –60°), mirror reversals (180°), and default reaches directly to the target (0°). The same multimodal structure appeared at the generalization target (Figure 2D, bottom; Best BIC: inner = −5851.3 with 7 modes; outer = −5432.0 with 6 modes; full mode breakdown in Table S2). These multimodal error distributions were again incompatible with the gradual error reduction account, which predicted errors confined between baseline (0°) and goal (60°). Instead, it aligned with learners actively testing multiple action–outcome hypotheses.

Why, then, did an efficiency–flexibility trade-off emerge between the inner- and outer-target groups? We addressed this in two ways. First, motor variability was substantially higher in the outer-target group (main effect of Group: *F* = 26.9, *p* <.001, *η*^2^ = 0.09; early adaptation post-hoc t-test: *t* = 5.6, *p* <.001), suggesting that the constrained geometry prompted greater exploration. Second, the groups differed strikingly in their movement-angle distributions during training (Figure 2D-E; JSD = 0.06, *p* =.003): the outer-target group explored a wider hypothesis space—showing more sign flips (–60°), more mirror reversals (180°), a stronger uniform component, and fewer reaches to the correct solution (60°). At the generalization target, this pattern reversed (JSD = 0.13, *p* <.001): the outer-target group produced fewer sign flips and more movements near the correct 60° rule.

Together, these results provide a process-level explanation for the efficiency–flexibility trade-off: the outer-target group explored more broadly—slower to discover a successful solution but ultimately generating a more flexible strategy. Importantly, the coexistence of a group-level trade-off with multimodal exploratory movements at the individual level is incompatible with the gradual error reduction account. Instead, these results support hypothesis testing as a novel process governing human sensorimotor learning.

### Experiment 2: Reproducing the Efficiency–Flexibility Trade-off Under Stringent Conditions

Experiment 1 revealed an efficiency–flexibility trade-off shaped by the spatial geometry of the training targets—a pattern consistent only with the hypothesis testing account. However, one methodological issue tempers this interpretation: the inner- and outer-target groups differed not only in target geometry but also in their spatial distance from the generalization target. Because generalization declines with distance (Heald et al., 2021, 2023; McDougle & Taylor, 2019), the outer-target group’s broader generalization could simply reflect that their generalization target was closer to the trained locations (110° in the outer vs. 170° in the inner condition).

To address this, we designed Experiment 2 (N = 280), again manipulating the level of environmental constraint via target geometry while equating the spatial distance between training and generalization targets (110° for both groups; Figure 1D). Participants in both groups adapted to *eight* training targets, allowing us to vary geometric structure while holding generalization distance constant. In the outer-target (constrained) group, the target geometry provided stronger cues that prompted the imposed rotation hypothesis. In contrast, the inner-target (unconstrained) group—marked by fewer outer targets and more inner targets—rendered multiple hypotheses plausible.

We reproduced the efficiency–flexibility trade-off under these stringent conditions. Inner group learned faster during early adaptation (Inner: 33.8° ± 3.1; Outer: 27.4° ± 4.0; *p* =.012). Both groups successfully compensated for the perturbation at the training targets (late adaptation: inner 52.2° ± 1.9.; outer 52.1° ± 2.0; *p* =.65), and neither exhibited implicit aftereffects (*p* <.001), confirming that performance reflected primarily strategic learning processes. Although the outer-target advantage was subtle, significant group differences still emerged (Figure 3A, left; cycles 13–16: Δμ = −3.0°, *d*_*circ*_ = 0.0, *p*_*perm*_ =.006; cycles 27–32: Δμ = −2.1°, *d*_*circ*_ = −0.1, *p*_*perm*_ =.010).

**Figure 3.**
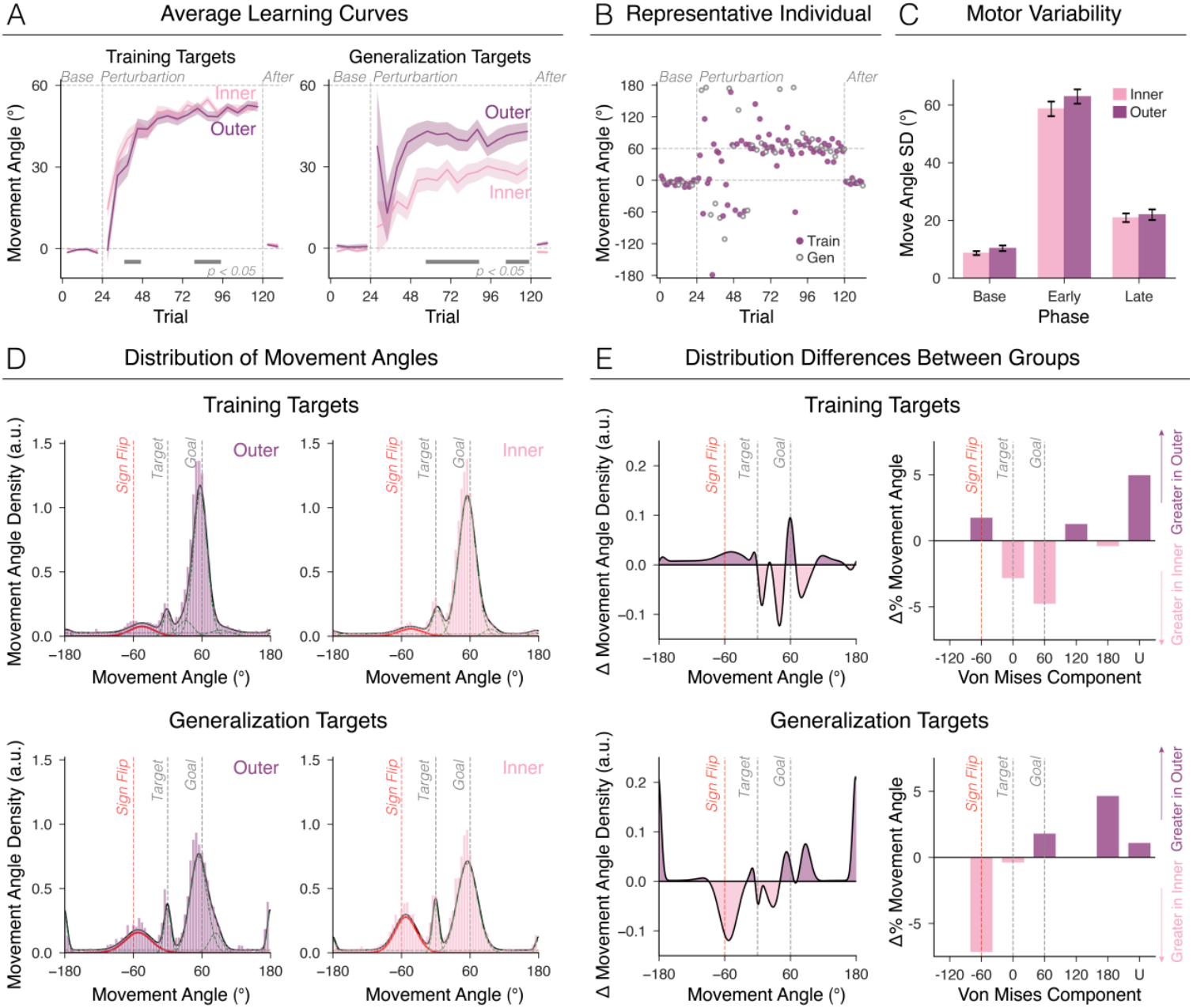
Reproducing the Efficiency–Flexibility Trade-off in Strategic Motor Adaptation under Stringent Conditions. **(A)** Learning curves for training **(left)** and generalization targets **(right)** in Experiment 2. Mean movement angle (°) is plotted for the outer-target (dark green) and inner-target (light green) groups; shaded regions denote SEM. Vertical dashed lines indicate perturbation onset and offset. Horizontal gray bars mark significant group differences identified via cluster-based permutation tests (p <.05). **(B)** Representative learner from the outer group. Filled purple dots correspond to movements to the training targets; open gray dots correspond to movements to the generalization targets. **(C)** SD of within-subject training-target movement angles across phases. “Base” = baseline, “Early” = early adaptation, “Late” = late adaptation. Error bars indicate SEM. **(D)** Movement angle distributions for training (top) and generalization (bottom) targets. Solid black lines show von Mises mixture fits; dashed red lines mark individual mixture components; histograms are rescaled to match the density range. Vertical dotted lines denote the training target (0°) and the required rotation solution (60°). Sign flips (–60°)—correct-magnitude reaches in the opposite/incorrect direction—are highlighted in red for illustration. **(E)** Distributional differences of von Mises fit (density) between inner- and outer-target groups for training and generalization targets **(left)**. Negative values indicate greater density in the inner-target group; positive values indicate greater density in the outer-target group. These differences are further illustrated by comparing the von Mises component weights (binned in 60° increments) and the uniform component **(right)**. Critically, outer-target learners exhibit greater exploration at the training targets (i.e., greater sampling of non-solution regions), a pattern that reverses at the generalization targets, consistent with an efficiency–flexibility trade-off.

Crucially, this pattern reversed at the generalization target (Figure 3A, right; interaction between Target Geometry x Target Type: F=6.3, p=0.012, η^2^=0.02): the inner-target group generalized less than the outer-target group. Significant group differences were observed throughout cycles 13–30 (Δμ = 14.9°, d_circ_ = 0.2, *p*_*perm*_ <.001) and cycles 35-40 (Δμ = 13.8°, *d*_*circ*_ = 0.2, *p*_*perm*_ =.001). Together, an efficiency–flexibility trade-off persisted even when generalization distance was equated.

Individual learning curves showed pronounced early motor variability (Figure 3C: Inner: 58.7° ± 2.5, Outer: 62.9° ± 2.5), which declined systematically over time (Late SD: Inner: 20.9° ± 1.5, Outer: 22.0° ± 1.8). These values were significantly larger than baseline sensorimotor noise (main effect of Phase: *F* = 829.8, *p* <.001, *η*^2^ = 0.75), consistent with participants sampling and refining multiple action–outcome hypotheses before converging on a successful strategy. Additionally, movement distributions were multimodal (Figure 3D, top; Best BIC: inner = −8729.6 with 6 modes; outer = −9382.1 with 7 modes; von Mises parameters in Table S3), with several modes far from the 60° solution. This same multimodal structure appeared at the generalization target (Figure 3D, bottom; Best BIC: inner = −5762.4 with 5 modes; outer = −5844.5 with 6 modes), consistent with learners testing hypotheses rather than gradually reducing error.

We then asked whether the two groups differed in how they explored—that is, in the hypotheses they sampled. Unlike Experiment 1, distributional differences in movement angles between groups were small (Figure 3D; training target: JSD = 0.03, *p* =.694; generalization target: JSD = 0.07, *p* =.088), and motor variability did not differ significantly between groups (Figure 3B; *F* = 2.33, *p* =.12, *η*^2^ = 0.008; post-hoc t-test: *t* = 1.2, *p* =.23)—an issue we revisit with a more sensitive cross-experiment analysis. Nonetheless, Experiment 2 replicated the pattern from Experiment 1: At the training targets, the outer-target group showed greater exploration— marked by larger uniform, sign flip (−60°), and mirror reversal (180°) components. Conversely, at the generalization target, the outer-target group was more likely to identify the correct 60° solution (Figure 3E).

Taken together, the efficiency–flexibility trade-off and the rich, multimodal exploratory patterns at the individual level provide convergent evidence that strategic motor learning is driven by hypothesis testing, not gradual error reduction.

### Combined analysis across experiments: The efficiency–flexibility trade-off emerges from a continuous shift in the mixture of individual strategies

While the efficiency–flexibility trade-off emerged in both experiments, it was pronounced in Experiment 1 and more subtle in Experiment 2. This difference likely reflects the weaker geometric contrast in the eight-target conditions: In Experiment 1, the two-target geometries differed completely in target placement, whereas in Experiment 2, the eight-target geometries substantially overlapped and differed only in the arrangement of the inner targets.

Across experiments an intriguing observational trend emerged: the ratio of inner (unconstrained) to outer (constrained) targets appeared to determine where each condition fell along the efficiency–flexibility continuum (2 outer → 8 outer → 8 inner → 2 inner). Pure inner-target environments (2-inner) provided the most unconstrained setting—supporting rapid learning but yielding narrow generalization—whereas pure outer-target environments (2-outer) imposed strong constraints, producing slower learning but broad generalization. The mixed eight-target conditions appeared to fall between these extremes, consistent with the idea that the trade-off scales in a graded manner with the proportion of inner versus outer targets.

We statistically demonstrate this continuum across experiments in several ways (N = 560). First, at the training targets, early adaptation increased continuously from 2-outer to 8-outer to 8-inner to 2-inner—precisely mirroring the proportion of outer (constrained) versus inner (unconstrained) targets (Figure 4A; mixed-effects regression: β = 6.0, *p* =.005). At the generalization target, this pattern reversed: early adaptation decreased in the same graded sequence (mixed-effects regression: β = –5.7, *p* <.001).

Second, exploratory behavior showed the same continuum: Summing the von Mises weights for non-solution (non–60°) modes—a proxy for exploration—revealed a graded increase from 2-inner to 2-outer at the training targets, mirroring the ratio of constrained outer targets; at the generalization target, this pattern reversed (Figure 4B). This again supports the idea that the efficiency–flexibility trade-off scales continuously with the ratio of constrained outer to unconstrained inner targets.

Third, we examined individual learning curves using an unsupervised machine-learning approach, clustering participants with soft dynamic time warping (see Methods). We restricted this analysis to participants who successfully learned during training (N = 476 out of 560; 85% learner rate), a conservative approach to ensure that clusters reflected genuine strategic differences rather than inattention or task disengagement. This unsupervised approach is enabled by our large online sample, revealing fine-grained learning profiles that would be difficult to detect in smaller in-person studies.

**Figure 4.**
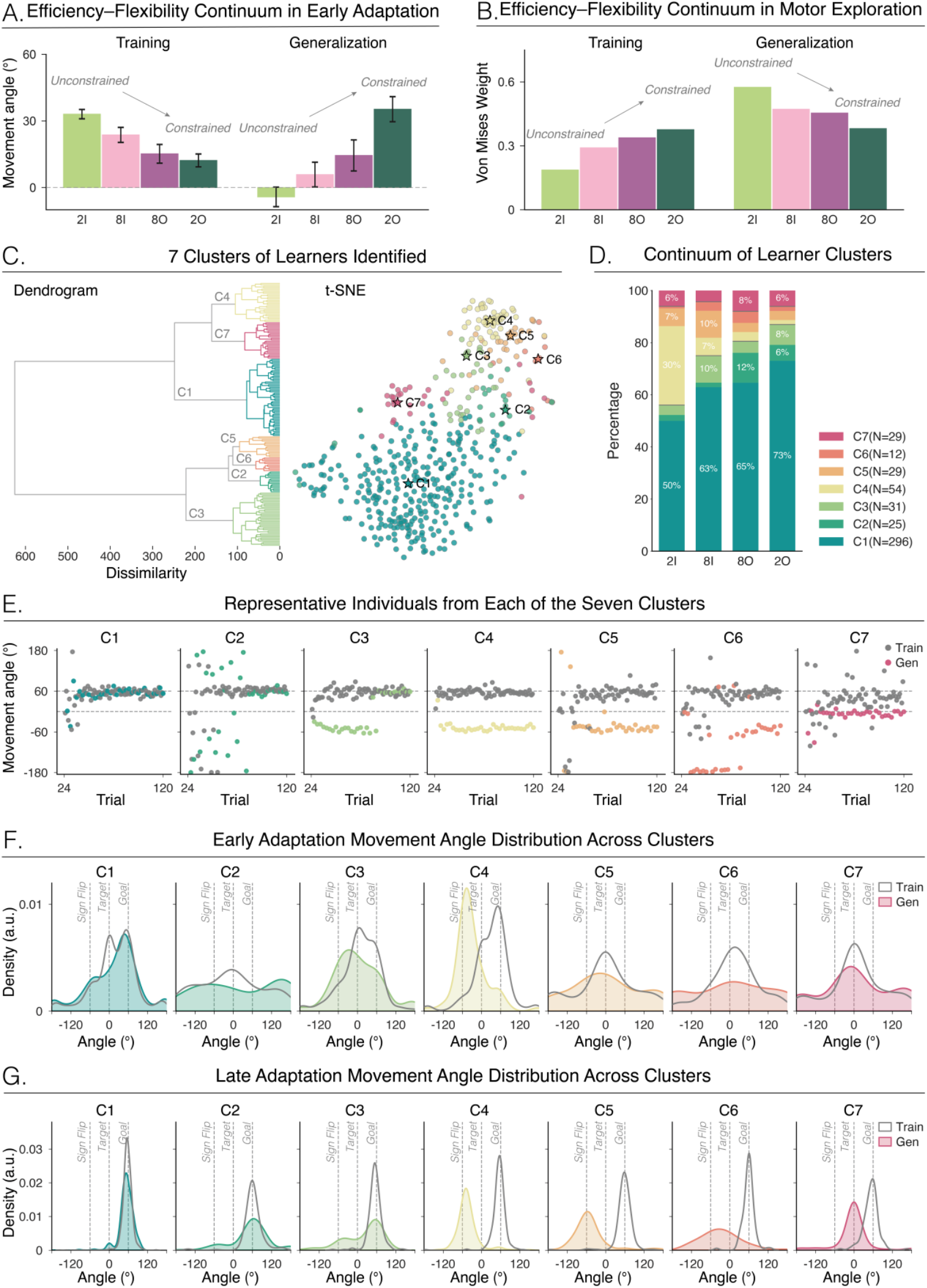
The efficiency–flexibility trade-off emerges from a continuous shift in the mixture of individual strategies. **(A)** Mean movement angle during early adaptation across the four experimental conditions: Experiment 1 (2-inner, 2-outer) and Experiment 2 (8-inner, 8-outer). Error bars indicate ± SEM across participants. **(B)** Total von Mises mixture weight at the non-solution (60°) locations across the four experimental conditions, indexing the degree of motor exploration across groups. **(C)** Dendrogram and t-SNE embedding showing learner clusters identified via pairwise distances computed using Soft Dynamic Time Warping. Colors denote cluster membership (C1–C7). **(D)** Cluster distribution as a function of target geometry. Stacked bar plots show the proportion of participants in each cluster across the four target-geometry conditions. Percentages < 5% are omitted for clarity. **(E)** Representative learning trajectories for each cluster, marked by a star in panel C. Gray dots indicate training trials; colored dots denote generalization performance. **(F, G)** Movement-angle distributions during early **(F)** and late **(G)** adaptation across the seven learner clusters. Solid lines with shade and dashed lines represent the distribution of movement angles at training and generalization target locations, respectively.

We observed seven distinct profiles (Figure 4C & 4E, C1–C7). Although all clusters achieved successful performance at the training target, they diverged markedly at the generalization target: some converged on the correct solution through distinct exploratory trajectories (C1: 62.2% of learners; C2: 5.3%; C3: 6.5%), whereas others never reached the correct strategy— settling instead on sign-flip solutions (C4: 11.3%; C5: 6.1%; C6: 2.5%) or defaulting to straight-to-target movements (C7: 6.1%). These qualitative differences were echoed in the distinct movement-angle distributions observed during early and late adaptation (Figure 4F–G), illuminating the rich and diverse hypothesis-testing behavior that defines strategic adaptation.

Critically, the efficiency–flexibility trade-off emerged from a continuous shift in the mixture of strategies (Figure 4D; *χ*^2^(18) = 75.8, *p* <.001): Groups with different target geometries comprised mixtures of exploratory strategies whose proportions shifted continuously with the ratio of outer to inner targets (2 outer → 8 outer → 8 inner → 2 inner). Outer-target geometries produced a higher proportion of learners who rapidly converged on the correct rotational rule (C1). Conversely, inner-target geometries yielded more participants in clusters that either failed to reach the correct solution (C4–C7) or required extended exploration before finding it (C2– C3). Together, these rich individual strategies, coupled with the continuum in the efficiency– flexibility trade-off shaped by environmental constraint, underscore hypothesis testing as a core process governing strategic motor adaptation.

## Discussion

### Summary of significant and novel findings

How do people discover an effective movement strategy when the environment changes—such as when using the trackpad on an unfamiliar laptop? Dominant theories of motor learning treat strategic adaptation as a form of passive gradient descent, in which the system incrementally reduces movement error. In contrast, we propose a novel process in which strategic adaptation operates through active hypothesis testing: learners generate candidate action–outcome hypotheses about the change in environment, evaluate them against sensory feedback, and iteratively refine both their actions and their beliefs.

Our results support the hypothesis testing framework. Using a visuomotor rotation task that isolates strategic motor adaptation, we found that constrained environments—defined by target geometry—slowed learning at the trained targets yet promoted generalization at untrained targets, with participants more often applying the correct rotational rule. In contrast, unconstrained environments accelerated learning but impaired generalization, with many participants failing to discover the correct rule and instead producing mirror reversals, sign flips, or default reaches to the target. This efficiency–flexibility trade-off at the group level is uniquely predicted by the hypothesis testing account.

The hypothesis testing account is further supported in several ways. First, analyses of individual learning curves reveal substantial exploratory variability—far exceeding the levels of sensorimotor noise predicted by the gradual error account. Second, movement-angle distributions were strikingly multimodal, exhibiting 6–7 peaks, whereas gradual error reduction predicts movements constrained between baseline and solution states. Third, learners showed remarkable diversity in the temporal dynamics of strategy discovery: some discovered the 60° (clockwise or counterclockwise) rotation immediately, others exhibited a discrete moment of insight, and still others explored extensively yet never discovered the correct rule. Together, hypothesis testing qualitatively captures the diversity of exploratory behavior, whereas the gradual error reduction account cannot.

### Consideration of alternative explanations

One recent proposal argues that strategic adaptation occurs through a discrete moment of insight, in which behavior jumps abruptly from baseline to the correct solution with no intervening exploration (Townsend et al., 2025). However, our data do not support this account. Only a small subset of individuals exhibited such abrupt ‘aha’ moments, whereas most participants showed systematic exploratory errors. Moreover, although insight models can *describe* the form of behavioral change, they cannot *explain* why such insights arise. In contrast, hypothesis testing specifies a putative psychological process that gives rise to insights: A subset of learners may entertain a small set of candidate hypotheses— “no perturbation” and the 60° rotation—and begin with a strong prior favoring no perturbation. As discrepant sensory evidence builds, the prior eventually flips; once a decision threshold is crossed, learners infer a rotated environment and abruptly re-aim (Kaplan & Simon, 1990; Becker et al., 2025).

Another proposal argues that strategic adaptation reflects a reinforcement-learning process (Darshan et al., 2014; Taylor & Ivry, 2011, 2012; Therrien et al., 2016; van Mastrigt et al., 2021, 2023; Velázquez-Vargas et al., 2024), in which successful actions are exploited and failures induce exploratory noise (also see: Cashaback et al., 2017). Although such models can generate early variability, they predict random, uniformly distributed error patterns—not the multimodal distributions observed in our data. Moreover, reinforcement-driven exploration typically unfolds in a slow and inefficient manner, akin to searching for a needle (the solution) in a haystack (the entire 360° workspace). This stands in contrast to the speed of learning observed here, who often converge on the solution within just a few trials (Uehara et al., 2019). Accordingly, our data are inconsistent with reinforcement-based accounts of strategic adaptation.

### Relating hypothesis testing to the broader psychological literature

Our data prompt a reinterpretation of prior claims that motor errors during sensorimotor learning—such as sign flips—reflect attentional lapses in *implementing* an already learned strategy (McDougle & Taylor, 2019): Specifically, sign flips are often treated as moments when participants momentarily apply the correct rule with the wrong sign. However, if sign flips were lapses, they should occur uniformly across learning or even increase later in the task as cognitive fatigue sets in. Instead, we observe the opposite: sign flips—and related errors such as mirror reversals—are more common early in learning and diminish over time. Therefore, we interpret these errors as evidence that participants were actively testing different action–outcome hypotheses while *discovering* an optimal re-aiming strategy (see Supplemental Materials for vignettes from free-responses illustrating the diverse strategies participants reported).

While our results clarify how people initially discover a new movement strategy, future work could examine how this learned structure guides adaptation to new perturbations—a process known as structure learning (Bond & Taylor, 2017; Braun et al., 2010; Heald et al., 2023; Wolpert et al., 2011; Wolpert & Flanagan, 2010). Hypothesis testing predicts that once a rotational mapping is inferred, belief in this hypothesis is strengthened, thereby facilitating the acquisition of future perturbations with similar rotational structures (Tian et al., 2025). Conversely, such consolidation would impede learning of different geometric mappings, such as reflections or translations. Evaluating these predictions would clarify how hypothesis testing informs structural learning.

Our results situate strategic motor adaptation within the broader landscape of hypothesis-driven cognition. Across many domains, people learn by generating and evaluating structured hypotheses: in problem solving, they test and revise predictions about an opponent’s next move to guide their own (Cowley & Byrne, 2004; Newell & Simon, 1972); in intuitive physics, humans evaluate alternative action–outcome predictions to determine how objects will fall or collide, or how to wield an unfamiliar tool (Allen et al., 2020); and in concept learning, they test different categorical boundaries to uncover latent rules that group stimuli into meaningful categories (Anderson, 1982; Steyvers et al., 2003). Viewed through this lens, strategic adaptation is best understood as a form of ‘embodied problem solving’—a process driven not by passive error reduction, but by the active generation and testing of action-outcome hypotheses (see hypothesis testing applied to other domains: (Kemp & Tenenbaum, 2008; Klayman & Ha, 1989; Lee et al., 2016; Levine, 1975; Nosofsky et al., 1994; Restle, 1962).

However, it is important to note that we are not claiming that hypothesis testing provides a superior account across all regimes of motor learning. For example, the gradual error reduction framework—typically formalized as a state-space or Kalman filter model—offers an excellent description of the implicit processes that support gradual sensorimotor recalibration (Burge et al., 2008; Smith et al., 2006; Padmanabhan et al., 2025). Reinforcement learning effectively captures skill acquisition, as learners refine speed, precision, and timing through repetition (Haith, 2025; Seethapathi et al., 2024). Our aim is not to supplant these frameworks but to broaden the vocabulary of motor learning to include hypothesis testing—a process that likely complements (Miyamoto et al., 2020), and at times competes (Albert et al., 2022) with, these other processes (for a review, see Therrien & Wong, 2022).

Our findings also contribute to a growing literature highlighting the central role of the goal in sensorimotor learning (Leow et al., 2018; Molinaro & Collins, 2023; Taylor & Ivry, 2013; Tsay et al., 2022; Padmanabhan et al., 2025; Villavicencio et al., 2025). Prior work shows that target geometry shapes implicit adaptation. We extend this to strategic adaptation: target geometry can bias learners toward particular action–outcome hypotheses. Specifically, outer-target geometries, which tacitly emphasize angular relationships across the workspace, cue learners to interpret the perturbation as a rotation. In contrast, inner-target geometries, which provide weaker cues for an angular structure, allow multiple hypotheses to remain plausible. Beyond its theoretical implications, manipulating features of the goal may offer a principled route to guiding learners toward more effective sensorimotor strategies in clinical or coaching contexts (Leech et al., 2022; Roemmich & Bastian, 2018).

Finally, a central contribution of this work is the recognition that group averages can obscure the very learning processes we seek to understand. Instead, we argue that learning processes are best understood through a careful dissection of individual learning trajectories—an approach long embraced in other areas of learning research but rarely applied in motor learning (Gallistel et al., 2004; Gallistel, 2013; Mercado III, 2008; Braver et al., 2010; Donner & Hardy, 2015; Maggi et al., 2024). To make this possible, we developed a suite of tools novel to motor learning, enabling us to capture the diversity, richness, and temporal dynamics of individual behavior—and, in turn, to reveal hypothesis testing as a core mechanism governing sensorimotor learning.

